# Regeneration from three cellular sources and ectopic mini-retina formation upon neurotoxic retinal degeneration in *Xenopus*

**DOI:** 10.1101/2023.05.12.540545

**Authors:** Parain Karine, Albert Chesneau, Morgane Locker, Caroline Borday, Perron Muriel

**Affiliations:** Université Paris-Saclay, CNRS, Institut des Neurosciences Paris-Saclay, 91400, Saclay, France

**Keywords:** retina, photoreceptor degeneration, cobalt chloride, regeneration, Müller glia, ciliary marginal zone, retinal pigmented epithelium, *Xenopus laevis*, *Xenopus tropicalis*

## Abstract

Regenerative abilities are not evenly distributed across the animal kingdom. Interestingly, the underlying modalities are also highly variable, even among closely related species. In fish or amphibians, retinal repair can involve the mobilization of different cellular sources, including stem cells of the ciliary marginal zone (CMZ), retinal pigmented epithelial (RPE) cells, or Müller glia. The mechanisms that trigger the recruitment of one cell type over another remain elusive. To investigate whether the magnitude of retinal damage might influence the regeneration modality of the *Xenopus* retina, we developed a model based on cobalt chloride (CoCl_2_) intraocular injection, allowing for a dose-dependent control of cell death extent. Analyses in *Xenopus laevis* revealed that limited CoCl_2_-mediated neurotoxicity only triggers cone cell loss and results in a few Müller glia cells reentering the cell cycle, without affecting CMZ cell activity or recruiting RPE cells. Conversely, we found that severe CoCl_2_-induced retinal degeneration not only potentializes the proliferative response of Müller cells, but also enhances CMZ cell proliferation and, unexpectedly triggers an RPE reprogramming event. Although Müller glia could not regenerate cones under these conditions, both CMZ and RPE-derived proliferative cells could. Strikingly, RPE reprogrammed cells self-organized into an ectopic layered mini retina-like structure laid on top of the original retina. It is thus likely that the injury paradigm determines the awakening of different stem-like cell populations exhibiting distinct neurogenic capacities. Besides, we surprisingly found that *Xenopus tropicalis* also has the ability to recruit Müller cells and reprogram its RPE following CoCl_2_-induced damage, whereas only CMZ cell proliferation was reported in previously examined degenerative models. Altogether, these findings highlight the critical role of the injury paradigm and reveal that three cellular sources can be reactivated in the very same degenerative model.

## INTRODUCTION

Some species, such as fish or amphibians, are endowed with a remarkable ability to regenerate their retina in response to injury. In contrast, only very limited regeneration spontaneously occurs within the mammalian eye. The mechanisms underlying such differential potential are mostly unknown but actively investigated in order to foresee strategies able to stimulate retinal self-repair in patients. Beyond such variability in efficiency lies another enigma related to the multiplicity of regeneration modalities. Different cellular populations can be mobilized to replace lost cells, including stem cells of the ciliary marginal zone (CMZ), the retinal pigmented epithelium (RPE), or Müller glial cells (Ail and Perron, 2017; Chiba, 2014; Fischer et al., 2013; Hamon et al., 2016; Hidalgo et al., 2014). The way a retina regenerates likely depends on the considered species and/or on the applied injury paradigm. For instance, it was reported that retinectomized *Xenopus tropicalis* individuals replenish their damaged retina by enhancing CMZ stem/progenitor cell proliferation, while *Xenopus laevis* ones mainly use RPE cell reprogramming (Araki, 2007; Miyake and Araki, 2014; Yoshii et al., 2007). Following mechanical injury or targeted rod cell ablation in *X. laevis*, no RPE dedifferentiation has been reported. However, Müller glial cells come into play, as observed in fish or newly hatched chick (Fischer, 2005; García-García et al., 2020; Goldman, 2014): they reprogram towards a stem/progenitor state, proliferate and generate new neurons (Langhe et al., 2017; Parain et al., 2022). The mechanisms that govern the recruitment of one cellular source over another and the implementation of these different regenerative processes remain elusive. The ability of *Xenopus* to regenerate its retina from either CMZ, RPE, or Müller cells, depending on the injury paradigm, makes these amphibian species powerful model systems to address this issue.

We here aimed to investigate whether the regenerative modality is dependent on the magnitude of retinal damage. To do so, we developed a novel *Xenopus* model of retinal degeneration, using cobalt chloride (CoCl_2_) as a neurotoxic agent, whose dose can be adapted to trigger various extents of neuronal cell death. Studies in different animal models (rodent, monkey, zebrafish, or pig) indeed previously showed that CoCl_2_ either causes selective photoreceptor loss or widely affects all retinal neurons, depending on its concentration (Hara et al., 2006; Kuehn et al., 2017; Medrano et al., 2018; Shirai et al., 2016). Of note, the mechanisms responsible for cobalt toxicity are not fully understood but are believed to rely, at least in part, on its ability to elicit chemical hypoxia (Chang et al., 2016; Muñoz-Sánchez and Chánez-Cárdenas, 2019). As expected, injections of different doses of CoCl_2_ into the *Xenopus laevis* retina yielded a range of degenerative phenotypes of increasing severity. The lower tested dose resulted in damage limited to cones. This triggered Müller glia reactivation but did not lead to any cone regeneration. With higher concentrations of CoCl_2_, cell losses were broader, extending to multiple retinal cell types. In these conditions, much more Müller cells re-entered into the cell cycle, and CMZ cell proliferation was found to be enhanced as well. Furthermore, and unexpectedly, such massive degeneration also induced RPE reprogramming, leading to the generation of a small newly formed retina facing the original one. Interestingly, in these conditions, new cones could be produced by CMZ and RPE cells but still not by Müller glia. Finally, this model of CoCl_2_-induced degeneration also revealed that *Xenopus tropicalis*, previously suspected to use mainly its CMZ (Miyake and Araki, 2014; Parain et al., 2022), is also able to efficiently reactivate its Müller cells and undergo RPE reprogramming. This work thus highlights that three cellular sources with contrasting regenerative capacities can be jointly recruited in both *X. laevis* and *X. tropicalis*, provided a specific degenerative context.

## RESULTS

### CoCl_2_ induces dose-dependent retinal degeneration in *Xenopus laevis*

To investigate whether CoCl_2_ intraocular injection could trigger retinal degeneration in *X. laevis* tadpoles, we tested several concentrations ranging from 10 to 100 mM. We next assessed cell death by TUNEL assay 2 days later (Fig. 1A-C). At the lowest dose, most TUNEL-positive cells were found in the photoreceptor layer. Higher CoCl_2_ concentrations led to the appearance of dying cells in the inner nuclear layer (INL) as well, and, ultimately, in the whole retina. In accordance with these apoptotic effects, retinal histology was also dose-dependently altered (Fig. 1D). A 10 mM CoCl_2_ injection did not seem to affect the retinal morphology, except for the length of the outer segments (OS), which appeared shortened, likely reflecting photoreceptor cell degeneration. In line with this, from 25 mM onwards, OS detached from the neural retina, and from 50 mM onwards, the outer nuclear layer (ONL) was no more distinguishable. Other retinal layers did not show major structural alterations at 25 mM, while the inner nuclear layer was found disorganized at higher concentrations. With 86% of dying cells localized in the ONL upon 10 mM CoCl_2_ injection (Fig. 1E), it is likely that this context of moderate neurotoxicity primarily affects photoreceptors. We thus decided to further characterize the low-dose effects. To determine the degeneration kinetics, a time course analysis was carried out from 1 to 6 days following injection (Fig. 1F). The number of TUNEL-positive cells was already significantly increased 1 day post-injection (dpi) compared to controls, and culminated at 2 dpi (Fig. 1G). At this time point, around half of photoreceptors were lost (data not shown). By 6 dpi, the number of apoptotic cells almost returned to a basal level, although it was still significantly higher than in control retinas. Together, these experiments reveal that a low dose of CoCl_2_ induces selective photoreceptor cell death in *X. laevis* during the next few days following intraocular injection.

**Figure 1.**
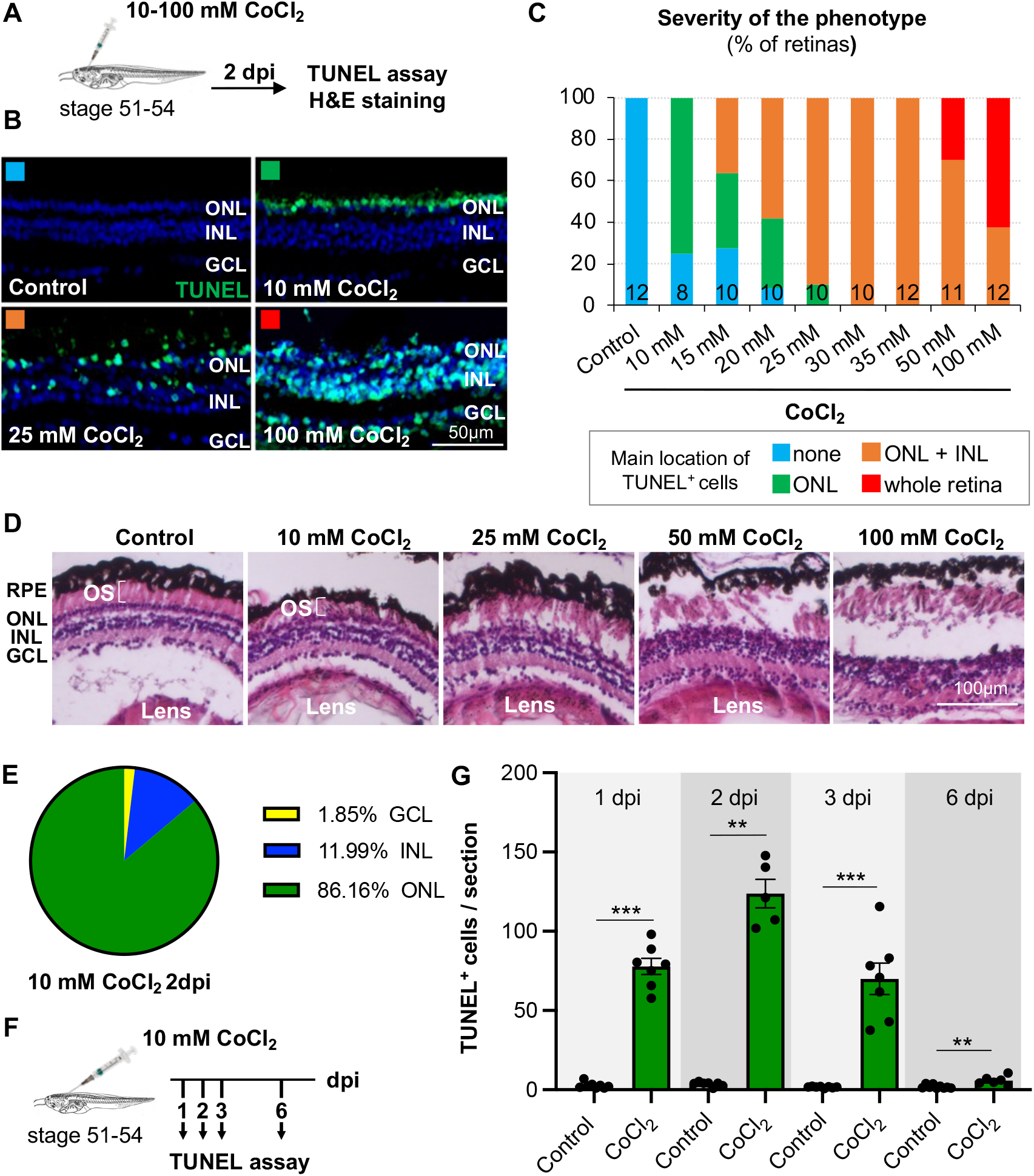
Dose-dependent effects of CoCl_2_ injections on retinal degeneration in *Xenopus laevis*. **(A)** Outline of the experimental procedure used in (B-E). Stage 51-54 *Xenopus laevis* tadpoles were intraocularly injected with a saline solution (control) or with CoCl_2_, at concentrations ranging from 10 to 100 mM CoCl_2_. They were then processed 2 days post injection (dpi) for TUNEL assay or Hematoxylin-eosin (H&E) coloration on retinal sections. **(B, C)** Analysis of cell death by TUNEL staining. Shown in (B) are representative images of retinal sections from control or CoCl_2_-injected tadpoles. Nuclei were counterstained with Hoechst. The graph in (C) represents the percentage of retinas exhibiting either no or very few apoptotic cells, apoptotic cells predominantly present in the ONL, in both the ONL and INL, or in the three retinal layers. The number of analyzed retinas is indicated in each bar. **(D)** Representative images of H&E staining on retinal sections following injection of CoCl_2_ at the indicated concentrations. **(E)** Pie chart of TUNEL^+^ cell distribution within the different retinal layers, 2 days after 10 mM CoCl_2_ injection. **(F)** Outline of the experimental procedure used in (G). Stage 51-54 tadpoles were intraocularly injected with a saline solution (control) or with 10 mM CoCl_2_. They were then processed for TUNEL assay at 1, 2, 3 or 6 dpi. **(G)** Corresponding quantification. Data are represented as mean ± SEM, and each point represents one retina. ** p < 0.01; *** p < 0.001 (Mann–Whitney tests). GCL: Ganglion cell layer, INL: Inner nuclear layer, ONL: Outer nuclear layer, OS: Outer segments of photoreceptors.

### A low dose of CoCl_2_ induces specific cone cell death in *Xenopus laevis*

Since about half of photoreceptors are rapidly lost in response to 10 mM CoCl_2,_ we then hypothesized that either cones or rods might be preferentially affected, as their ratio is about 50:50 in *Xenopus*. We thus undertook immunolabelling of both cell types at 3, 7, and 14 dpi, using Rhodopsin and S/M opsin as specific markers for rods and cones, respectively (Fig. 2A). The overall Rhodopsin labelling did not appear to decrease compared to controls, but revealed rod OS abnormal morphology at 3 and 7 dpi, with segments being likely tilted rather than neatly arranged in parallel. This phenotype was no more evident by 14 dpi (Fig. 2B). These data suggest that rods are only transiently affected by a 10 mM CoCl_2_ treatment, and then eventually recover. In contrast, we found that cone OS labelling was highly disorganized as soon as 3 dpi, and then decreased with time until virtual disappearance by 14 dpi (Fig. 2C, D). Finally, double TUNEL and S/M opsin staining confirmed that apoptotic cells were predominantly cones (Fig. 2E). Overall, these results highlight that moderate CoCl_2_-mediated neurotoxicity can serve as an experimental paradigm for cone-specific degeneration in *X. laevis*.

**Figure 2.**
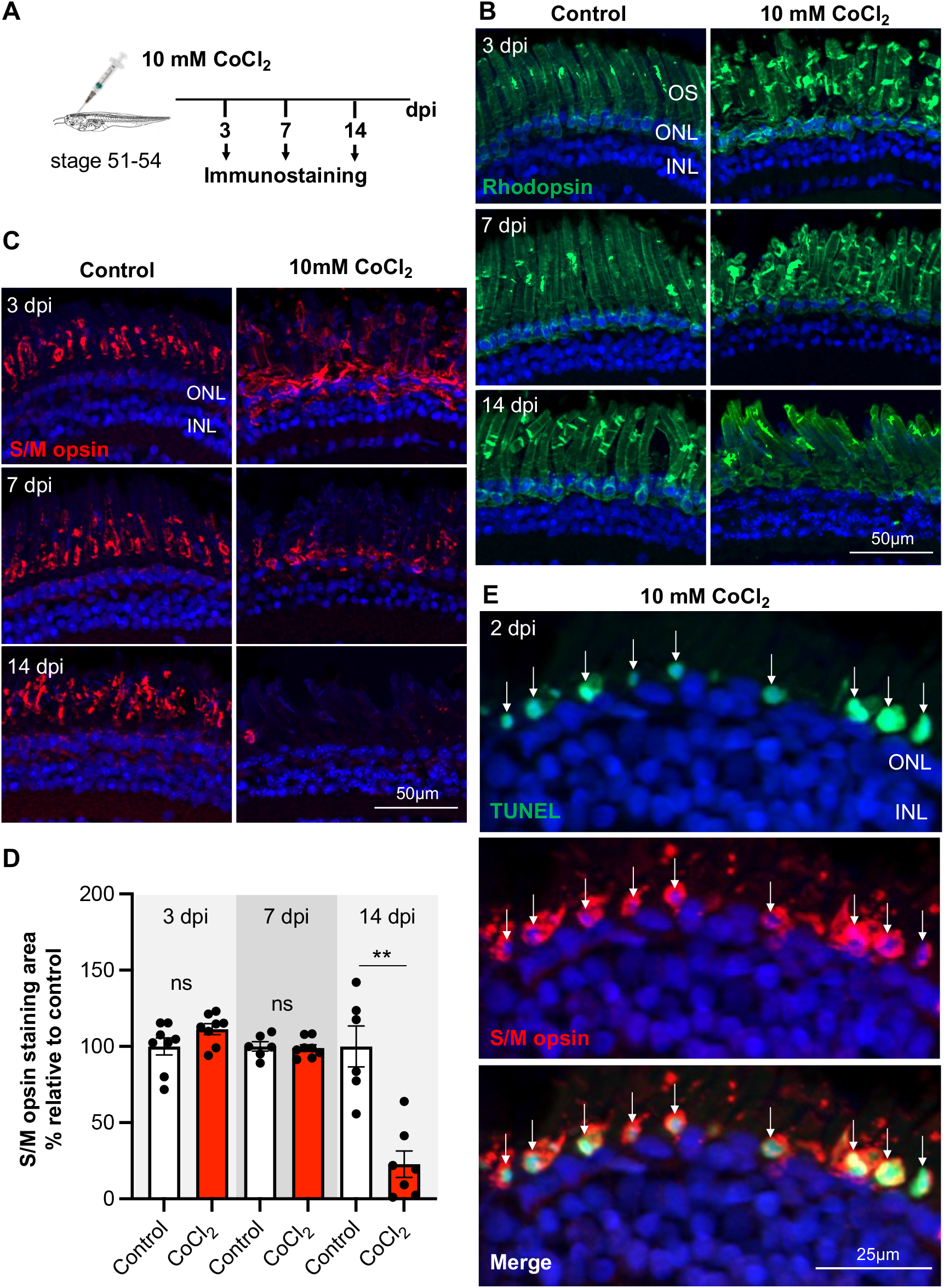
Specific cone cell death following a 10 mM CoCl_2_ intraocular injection in *Xenopus laevis*. **(A)** Outline of the experimental procedure used in (B-D). Stage 51-54 *Xenopus laevis* tadpoles were intraocularly injected with a saline solution (control) or with 10 mM CoCl_2_. They were then processed for immunostaining at 3, 7, or 14 dpi. **(B, C)** Representative images of S/M opsin (B) and Rhodopsin (C) staining. **(D)** Quantification of S/M opsin staining area relative to control. Data are represented as mean ± SEM, and each point represents one retina. ** p < 0.01, ns: non-significant (Mann–Whitney tests). **(E)** Representative images of retinal sections co-labelled for TUNEL and S/M opsin performed at 2 dpi in 10 mM CoCl_2_ injected-tadpole. Arrows point to double-positive cells. Cell nuclei are counterstained with Hoechst. INL: Inner nuclear layer, ONL: Outer nuclear layer, OS: Outer segments of photoreceptors.

### CoCl_2_-mediated cone degeneration induces Müller glial cell proliferation but does not trigger any cone regeneration in *Xenopus laevis*

We previously showed that conditional rod cell ablation leads to Müller cell proliferation and subsequent rod regeneration (Langhe et al., 2017). To assess whether Müller cells also respond to CoCl_2_-induced cone degeneration, we performed EdU incorporation assays in 10 mM CoCl_2_-injected tadpoles. EdU was made available for 3 days before fixation at either 3, 6, 14, or 21 dpi (Fig. 3A). The number of proliferating cells was significantly increased in CoCl_2_-injected retinas compared to controls from 6 dpi onwards, reaching a peak of 20 EdU-positive cells per section on average at 14 dpi (Fig. 3B, C). EdU-positive nuclei were predominantly found in the INL (Fig. 3C, D), suggesting that these cells may be Müller cells that have re-entered the cell cycle. To confirm their identity, we performed a double immunolabelling of PCNA (Proliferating Cell Nuclear Antigen), a marker of proliferative cells, and Yap (Yes-associated Protein), a marker of Müller glia (Hamon et al., 2017; Hamon et al., 2019; Rueda et al., 2019) (Fig. 3E, F). We found that 65% of PCNA-positive cells were expressing Yap. Of note, these double-labelled cells were observed both in the INL and the ONL, likely reflecting the migratory behaviour of reactive Müller cell nuclei. To examine whether these reactivated Müller cells can regenerate cones, we analyzed S/M opsin expression at 2 months post-injection (mpi) (Fig. 3G, H). A striking absence of staining was observed, suggesting the inability of *X. laevis* retinas to replace lost cones in this CoCl_2_ injury paradigm. In contrast, and in agreement with the aforementioned recovery of early rod defects (Fig. 2B), Rhodopsin staining appeared unaffected (Fig. 3H). Overall, these data suggest that selective cone cell death in *X. laevis* retinas promotes Müller glia cell cycle re-entry but is insufficient to trigger subsequent cone regeneration. Of note, considering the regenerative potential of the CMZ (Parain et al., 2022; Yoshii et al., 2007), we also investigated whether this region might respond as well. CMZ size, assessed following EdU labelling, was found similar in CoCl_2_-injected and control retinas at all time points examined, suggesting no enhancement of CMZ cell proliferation (Fig. S1). This indicates that targeted cone cell death in this CoCl_2_ injury paradigm does not trigger any regenerative response of retinal stem/progenitor cells.

**Figure 3.**
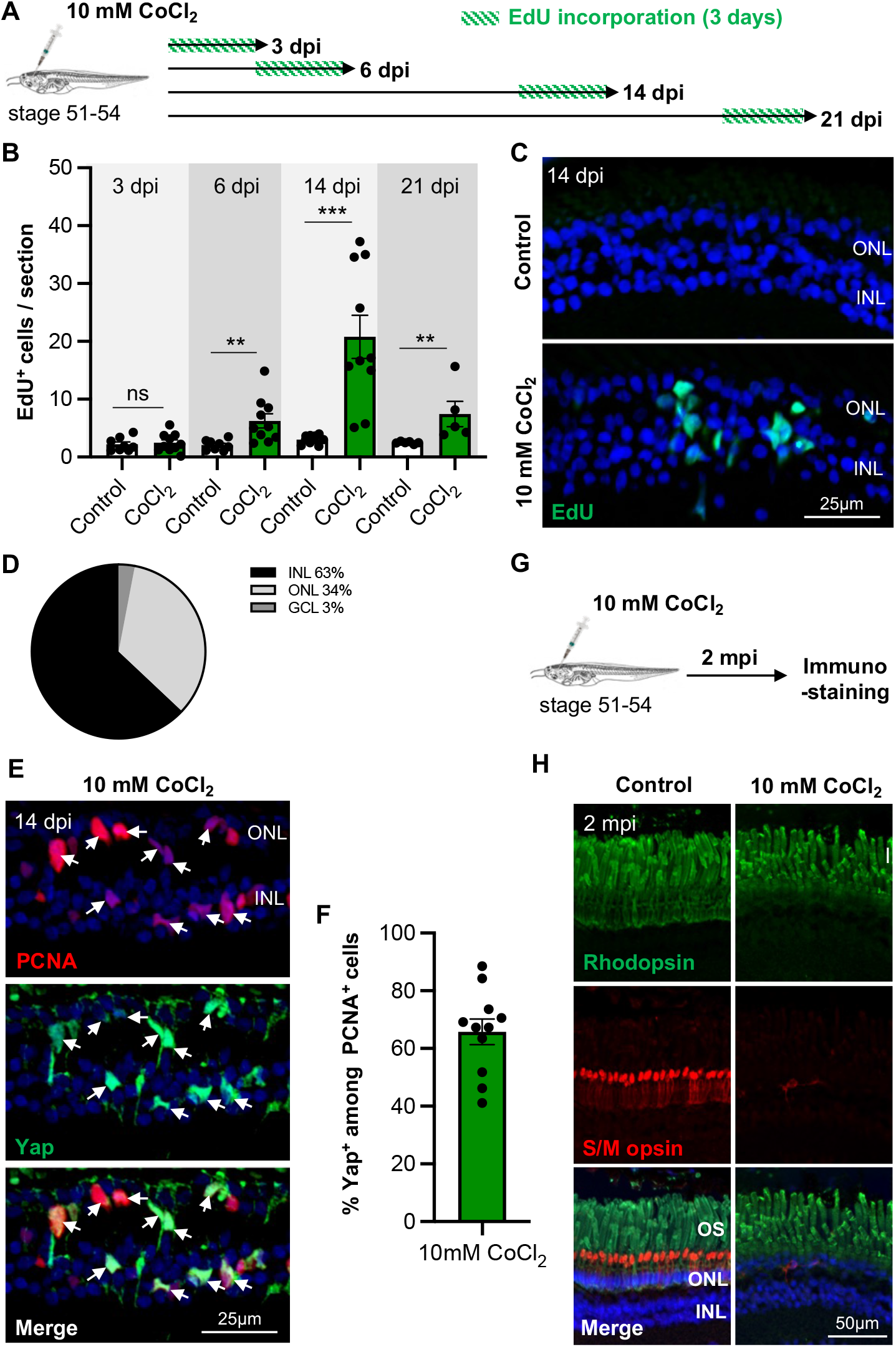
Müller glial cell proliferation without subsequent cone regeneration following a 10 mM CoCl_2_ intraocular injection in *Xenopus laevis*. **(A)** Outline of the experimental procedure used in (B-D). Stage 51-54 *Xenopus laevis* tadpoles were intraocularly injected with a saline solution (control) or with 10 mM CoCl_2_. They were then exposed for 3 days to EdU before being processed for analysis at 3, 6, 14, or 21 dpi. **(B)** Quantification of EdU-positive cells. **(C)** Representative images of EdU-labelled retinal sections at 14 dpi. **(D)** Pie chart of EdU-positive cell distribution within the different retinal layers at 14 dpi in CoCl_2_-injected tadpoles. **(E)** Representative images of retinal sections co-labelled for PCNA (proliferative cell marker) and Yap (Müller cell marker) at 14 dpi. Arrows point to double-positive cells. **(F)** Quantification of Yap^+^ cells among PCNA-labelled ones. **(G)** Outline of the experimental procedure used in (H). Stage 51-54 tadpoles were intraocularly injected with a saline solution (control) or with 10 mM CoCl_2_. They were then processed for immunostaining 2 months post-injection (mpi). **(H)** Representative images of retinal sections co-labelled for Rhodopsin and S/M-opsin. In (C, E, H), nuclei are counterstained with Hoechst. In graphs, data are represented as mean ± SEM, and each point represents one retina. ns: not significant, **p < 0.01, ***p < 0.001 (Mann-Whitney tests). INL: Inner nuclear layer, ONL: Outer nuclear layer, OS: Outer segments of photoreceptors.

### A high dose of CoCl_2_ induces bipolar and photoreceptor cell death in *Xenopus laevis*

We next hypothesized that the failure to regenerate new cone cells upon their selective loss was due to insufficient retinal damage among other retinal cell types. To investigate this question, we injected a higher dose of CoCl_2_ (25 mM) to trigger a broader degenerative context. At this concentration and as stated above (Fig. 1C), TUNEL-positive cells were found in both the INL and ONL, and the peak of apoptosis occurred between 2 to 4 dpi (Fig. 4A-C). To determine the identity of retinal cell types undergoing cell death, we first coupled the TUNEL assay with immunolabelling for Otx2, a marker of photoreceptor and bipolar cells. Double-labelled cells were observed in both the ONL and INL, suggesting that in addition to photoreceptors, some bipolar cells were also apoptotic (Fig. 4D). In line with this, the number of Otx2-positive cells was found decreased in both layers compared to the control situation (Fig. 4E, F). In contrast, analysis of Pax6-labelled cells did not reveal any difference between CoCl_2_-injected and control retinas (Fig. 4E, G). Pax6 being a marker of amacrine and ganglion cells, these cell types are thus likely spared from CoCl_2_-neurotoxic effects. Similarly, we did not detect any cell death in the RPE from 2 to 14 dpi in this injury paradigm (Fig. 4H and data not shown). Finally, we further examined photoreceptors by immunostaining. Cone OS showed aberrant morphology, and their swelling was associated with a temporary increase in S/M opsin labelling at 3 dpi. By 14 dpi, no more staining could be observed (Fig.4I, J). This suggests severe cone cell death, as observed with the 10 mM dose. Rods seemed less affected, although serious OS morphological defects were detectable from 3 dpi onwards. By 14 dpi, Rhodopsin staining was no longer homogeneous, and regions devoid of labelling could be observed (Fig. 4K). Overall, such 25 mM CoCl_2_ treatment affects bipolar cells and both rod and cone photoreceptors, with the most severe impact occurring on cones.

**Figure 4.**
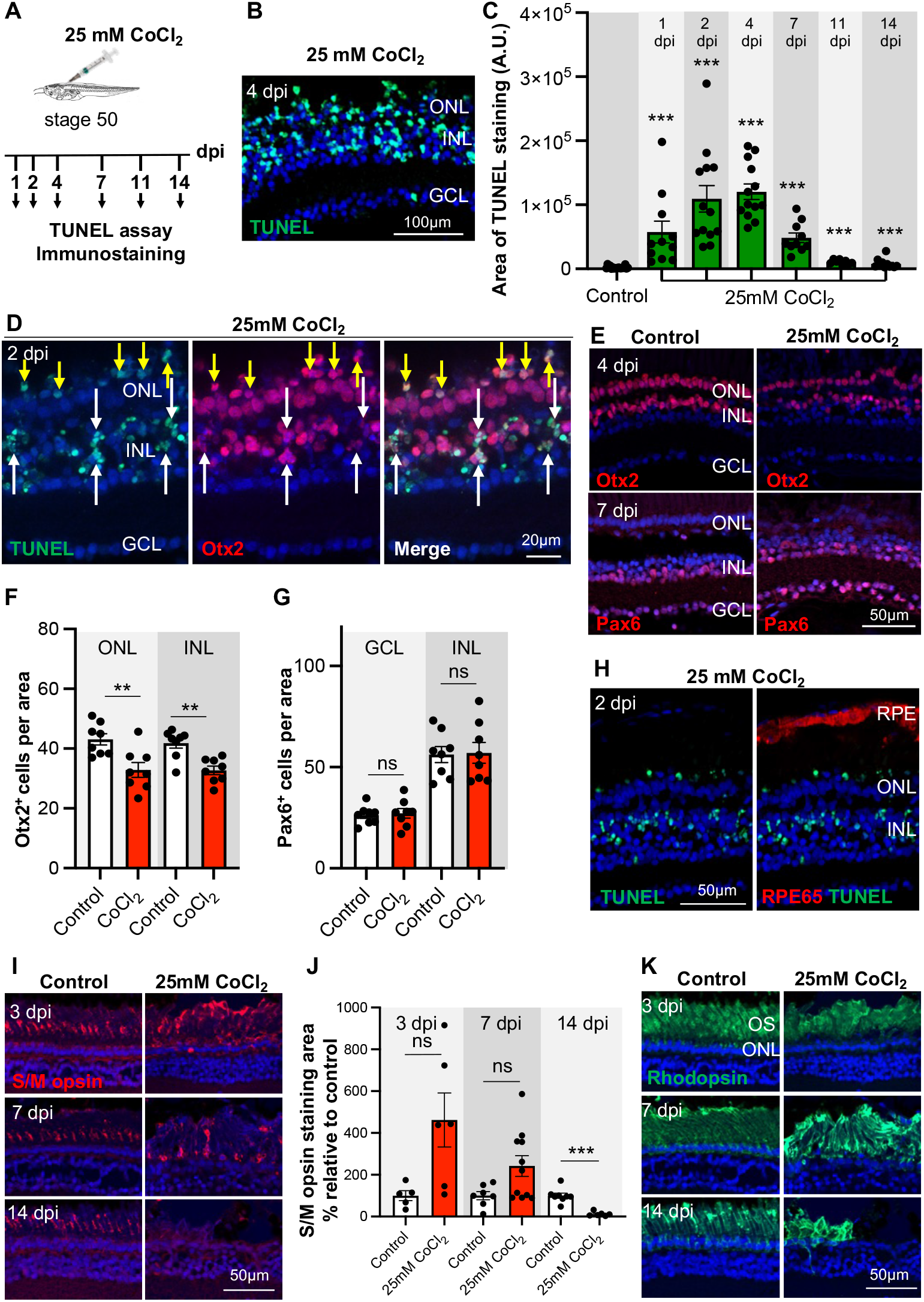
Bipolar and photoreceptor cell death following a 25 mM CoCl_2_ intraocular injection in *Xenopus laevis*. **(A)** Outline of the experimental procedure used in (B-K). Stage 51-54 *Xenopus laevis* tadpoles were intraocularly injected with a saline solution (control) or with 25 mM CoCl_2_. They were then processed for either TUNEL assay or immunostaining at different time points, as indicated. **(B, C)** Analysis of cell death by TUNEL staining. Shown in (B) is a representative section of CoCl_2_-injected retinas at 4 dpi. The graph in (C) shows the quantification of the TUNEL staining area from 1 to 14 dpi. **(D, H)** Representative images of retinal sections co-labelled for TUNEL and either Otx2 (a photoreceptor and bipolar cell marker) or RPE65 (a RPE marker) at 2 dpi. Arrows point to double-positive cells within the ONL (yellow) and INL (white). **(E-G)** Immunostaining analysis of Otx2 and Pax6 (an amacrine and ganglion cell marker) expression at 4 and 7 dpi, respectively. Representative retinal sections are shown in (E) and corresponding quantifications in (F, G). **(I-K)** Immunostaining analysis of S/M opsin and Rhodopsin expression on retinal sections at 3, 7, or 14 dpi. Shown in (J) is the quantification of S/M opsin staining area relative to control. In all images, nuclei are counterstained with Hoechst. In graphs, data are represented as mean ± SEM, and each point represents one retina. ns: not significant, **p < 0.01, ***p < 0.001 (Mann-Whitney tests). GCL: Ganglion cell layer, INL: Inner nuclear layer, ONL: Outer nuclear layer, RPE: Retinal pigmented epithelium.

### A high dose of CoCl_2_ triggers massive proliferation of Müller glia in *Xenopus laevis*

We next sought to investigate whether a broad degenerative context might more efficiently break Müller cell dormancy than a milder one. We indeed found that the amount of EdU- or PCNA-labelled cells was drastically enhanced upon injection of 25 mM CoCl_2_ compared to 10 mM (Fig. 5A-C). Nearly 80% of PCNA-positive cells were Müller glia, as assessed by their co-staining with Yap (Fig. 5D, E). Of note, a large amount of these proliferative Müller cells was found in the ONL. This was particularly striking at 14 dpi (Fig. 5D, F), indicating important nuclear migration events. As a complementary approach to confirm the identity of these numerous cycling cells, we established a transgenic reporter line, *Tg(her4:eGFP)*, that allows the visualization of Müller cell nuclei (Fig. 5G, H). Following injection of 25 mM CoCl_2_ in the eyes of *Tg(her4:eGFP)* tadpoles, we could indeed clearly see that the large majority of PCNA-positive nuclei were GFP-positive as well (Fig. 5I, J). Altogether, these results reveal that a 25 mM CoCl_2_ treatment leads to a massive Müller glia cell cycle re-entry, whereas the 10 mM dose showed much more limited effects. This suggests that either the magnitude of injury and/or the nature of dying cells influences the number of *X. laevis* Müller cells that exit quiescence.

**Figure 5.**
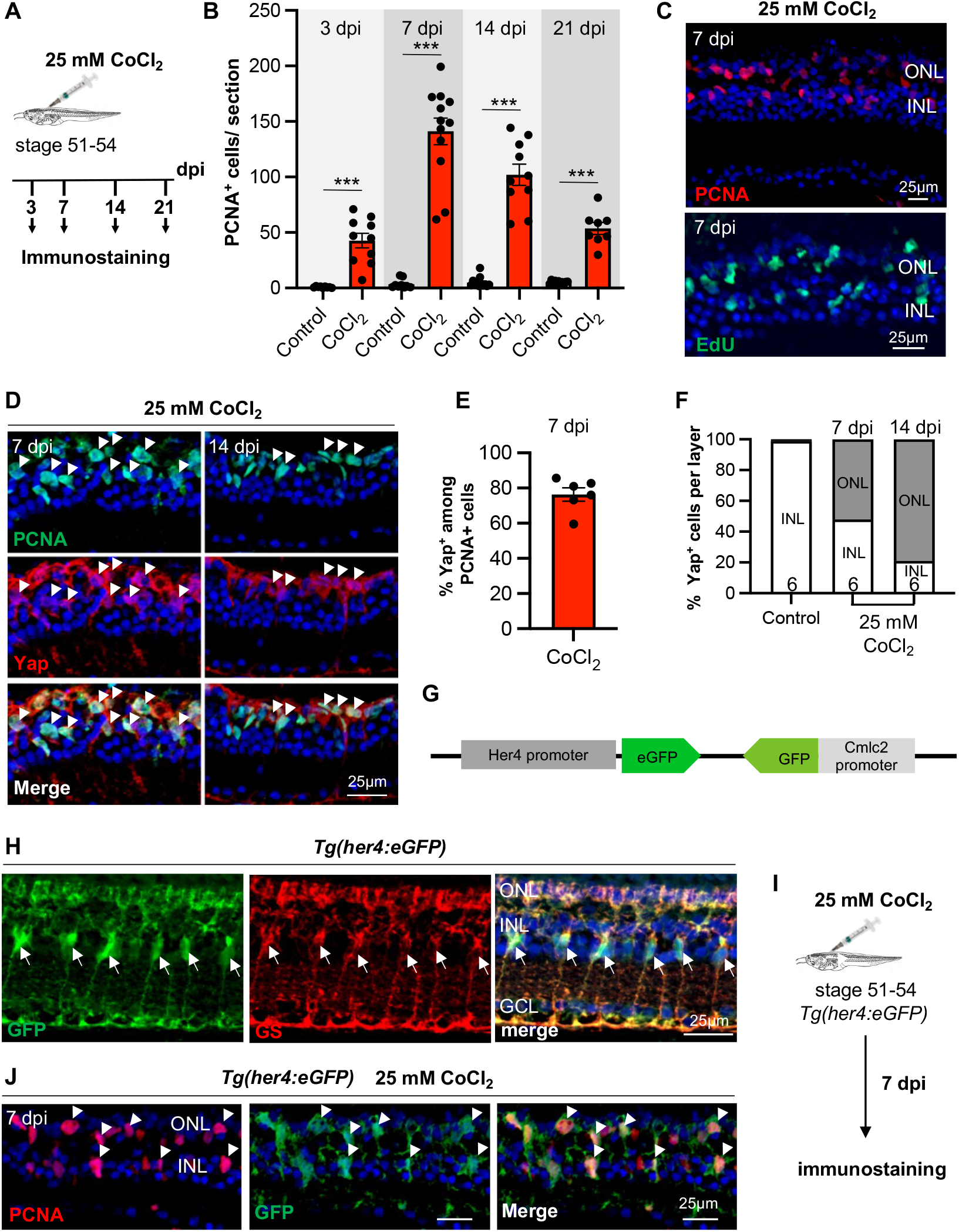
Massive Müller glia cell proliferation following a 25 mM CoCl_2_ intraocular injection in *Xenopus laevis*. **(A)** Outline of the experimental procedure used in (B-F). Stage 51-54 *Xenopus laevis* tadpoles were intraocularly injected with a saline solution (control) or with 25 mM CoCl_2_. They were then processed for immunostaining at different time points, as indicated. For EdU labelling, tadpoles were exposed to EdU three days prior to fixation. **(B, C)** Quantification of PCNA-positive cells and representative images of PCNA- or EdU-labelled retinal sections from CoCl_2_-injected tadpoles at 7 dpi. **(D-F)** Analysis of PCNA and Yap co-labelling at 7 and 14 dpi. Shown in (D) are representative images of retinal sections. Arrows point to double-positive cells. The graph in (E) is the quantification of Yap^+^ cells among PCNA-labelled ones. Shown in (F) is the distribution of Yap^+^ cells in the different retinal layers. **(G)** Schematic representation of the transgene *Tg(her4:eGFP)*. The *Her4* promoter drives *eGFP* expression in Müller glia, while the *Cmlc2* (cardiac myosin light chain 2) promoter drives it in the heart, allowing for transgenic tadpole screening. **(H)** Representative images of retinal sections from *Tg(her4:eGFP)* tadpoles, co-labelled for GFP and Glutamine Synthetase (GS; a marker of Müller cells). Arrows point to double-labelled cells. **(I)** Outline of the experimental procedure used in (J). Stage 51-54 transgenic *Tg(her4:eGFP)* tadpoles were intraocularly injected with a saline solution (control) or with 25 mM CoCl_2_. They were then processed for PCNA and GFP co-immunostaining at 7 dpi. **(J)** Representative images of retinal sections. Arrowheads point to double-positive nuclei. In all images, nuclei are counterstained with Hoechst. In (B, D), data are represented as mean ± SEM, and each point represents one retina. In (F), the number of analyzed retinas is indicated at the bottom of each bar. ***p < 0.001 (Mann-Whiney tests). GCL: Ganglion cell layer, INL: Inner nuclear layer, ONL: Outer nuclear layer.

### A high dose of CoCl_2_ enhances CMZ proliferative activity in *Xenopus laevis*

We next examined whether, in addition to Müller glia, CMZ cells might be mobilized as well in response to 25 mM CoCl_2_-induced degeneration. CMZ size, measured on PCNA-labelled sections, was found increased in CoCl_2_-injected retinas compared to control ones, from 3 dpi onwards and with a peak at 14 dpi (Fig. 6A, B). Thus, contrasting again with the 10 mM situation (Fig. S1), a broad degenerative context not only potentiates Müller cell recruitment, but also enhances the basal proliferative activity of CMZ cells.

**Figure 6.**
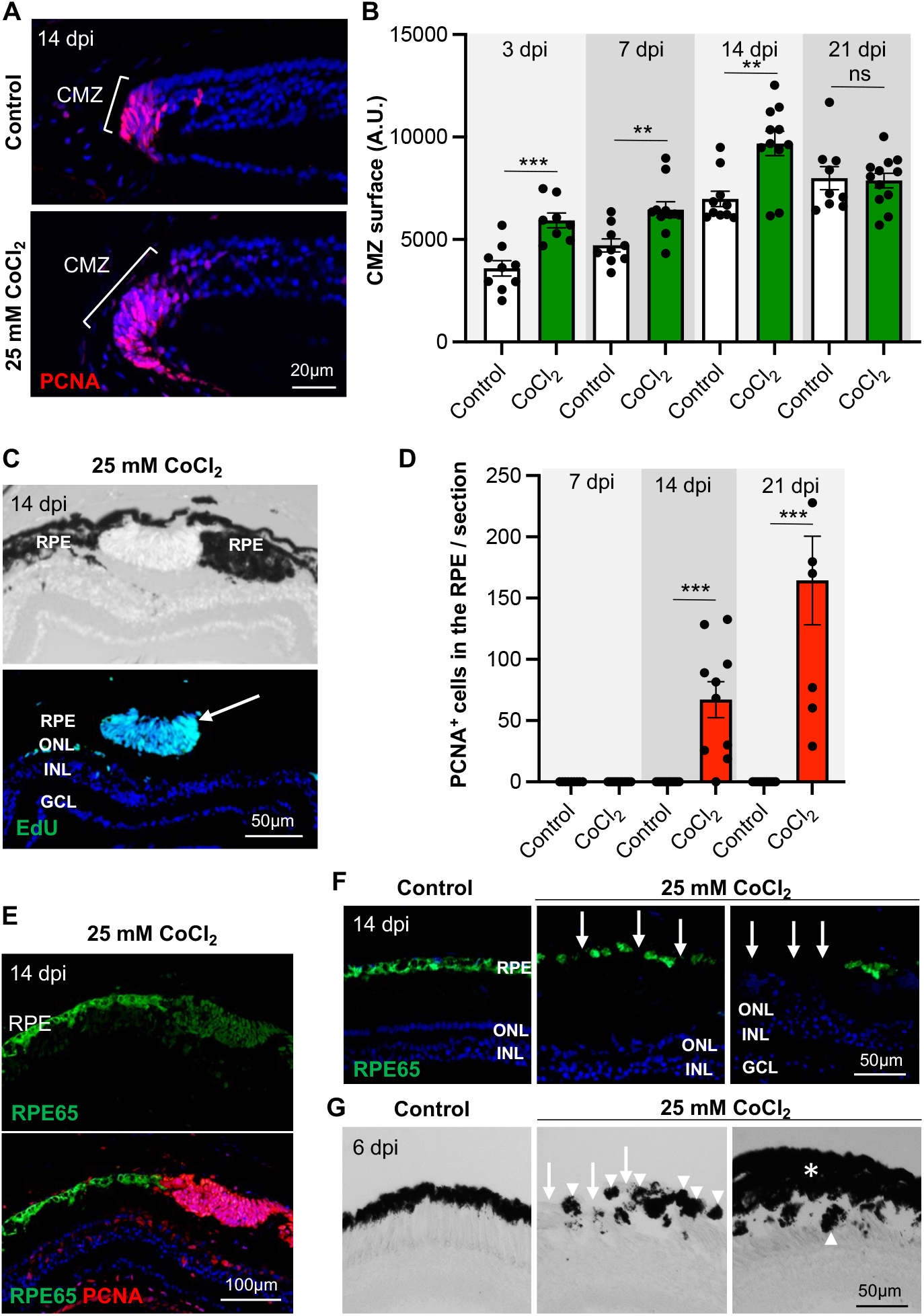
CMZ cell proliferation and RPE reprogramming following a 25 mM CoCl_2_ intraocular injection in *Xenopus laevis*. **(A, B)** Analysis of CMZ cell proliferation at different time points, as indicated, following injection of a saline solution (control) or 25mM CoCl_2_. Shown in (A) are representative images of PCNA staining within the CMZ. Shown in (B) is the quantification of the CMZ surface from 3 to 21 dpi. **(C)** Representative images (brightfield and EdU labelling) of retinal sections from stage 51-54 tadpoles, 14 days after injection of 25 mM CoCl_2_. Tadpoles were exposed to EdU for three days prior to fixation. The arrow points to a cluster of proliferative cells in the RPE layer. **(D)** Quantification of PCNA-positive cells within the RPE, 7, 14, and 21 days after injection of a saline solution (control) or 25 mM CoCl_2_. **(E)** Representative images of retinal sections from CoCl_2_-injected tadpoles, co-labelled for RPE65 and PCNA at 14 dpi. **(F, G)** RPE65 immunolabelling (F; 14 dpi) or brightfield images (G; 6 dpi), following injection of a saline solution (control) or 25 mM CoCl_2_. Arrows point to RPE defects (loss of pigmentation or RPE65 staining), arrowheads point to RPE pellets, and asterisks indicate RPE thickening. In all fluorescent images, nuclei are counterstained with Hoechst. In graphs, data are represented as mean ± SEM, and each point represents one retina. ns: not significant, **p < 0.01, ***p < 0.001 (Mann-Whitney tests). CMZ: Ciliary marginal zone, GCL: Ganglion cell layer, INL: Inner nuclear layer, ONL: Outer nuclear layer, RPE: Retinal pigmented epithelium.

### A high dose of CoCl_2_ triggers RPE reprogramming in *Xenopus laevis*

In addition to the aforementioned increased proliferation of Müller and CMZ cells, another phenotypic trait induced by 25 mM CoCl_2_ drew our attention. Bulge-like masses of non-pigmented cells were occasionally observed within the RPE layer. These cells incorporated EdU, showing their proliferative activity (Fig. 6C). Further PCNA labelling revealed their presence from 14 dpi onwards (Fig. 6D). The association of cell cycle re-entry and pigmentation loss is highly reminiscent of an RPE reprogramming event, as occurring following complete retinectomy (Yoshii et al., 2007). In line with this idea, these proliferative clusters expressed the RPE65 marker, confirming their RPE origin (Fig. 6E). The intensity of RPE65 staining was, however, decreased compared with adjacent non-proliferative RPE cells, suggesting an ongoing dedifferentiation process. Of note, the persistence of RPE65 expression at a lower level was previously reported during RPE reprogramming in adult retinectomized newts (Chiba et al., 2006). Besides, we found that the RPE structure was globally impaired, with discontinuities in RPE65 staining (Fig. 6F) and multiple pigmentation defects, such as loss of pigmentation or thickening of the pigmented layer (Fig. 6G). Altogether, these results suggest that in addition to causing broad alterations of RPE integrity, a 25 mM CoCl_2_ treatment also triggers cellular reprogramming in discrete regions of the RPE layer.

### A high dose of CoCl_2_ triggers CMZ-, but not Müller glia-dependent cone regeneration in *Xenopus laevis*

We next wished to investigate retinal cell repair following severe degeneration driven by CoCl_2_. We first performed an EdU pulse-chase experiment in *X. laevis*, in order to foresee whether the intense proliferative response of Müller cells might correlate with a powerful regenerative efficiency. However, and surprisingly, although many EdU-positive Müller cells were found in the ONL at 21 dpi and 1 mpi, only a very small fraction of them were co-labelled with a cone marker (Fig. 7A, B). In fact, almost no S/M opsin labelling could be detected in the central retina at 1 mpi (Fig. 7B), nor later at 2 mpi (Fig. 7C). Similarly, we identified only very few EdU-positive cells co-labelled with Rhodopsin (Fig. 7D). This indicates a virtual absence of Müller cell-dependent photoreceptor regeneration in this model. Most EdU-labelled cells within the INL were Otx2-positive, indicating a bipolar identity. Only rare EdU-labelled Pax6^+^ amacrine cells were found (Fig. 7D). This is consistent with bipolar cells being the main type of interneuron affected (Fig. 4F).

**Figure 7.**
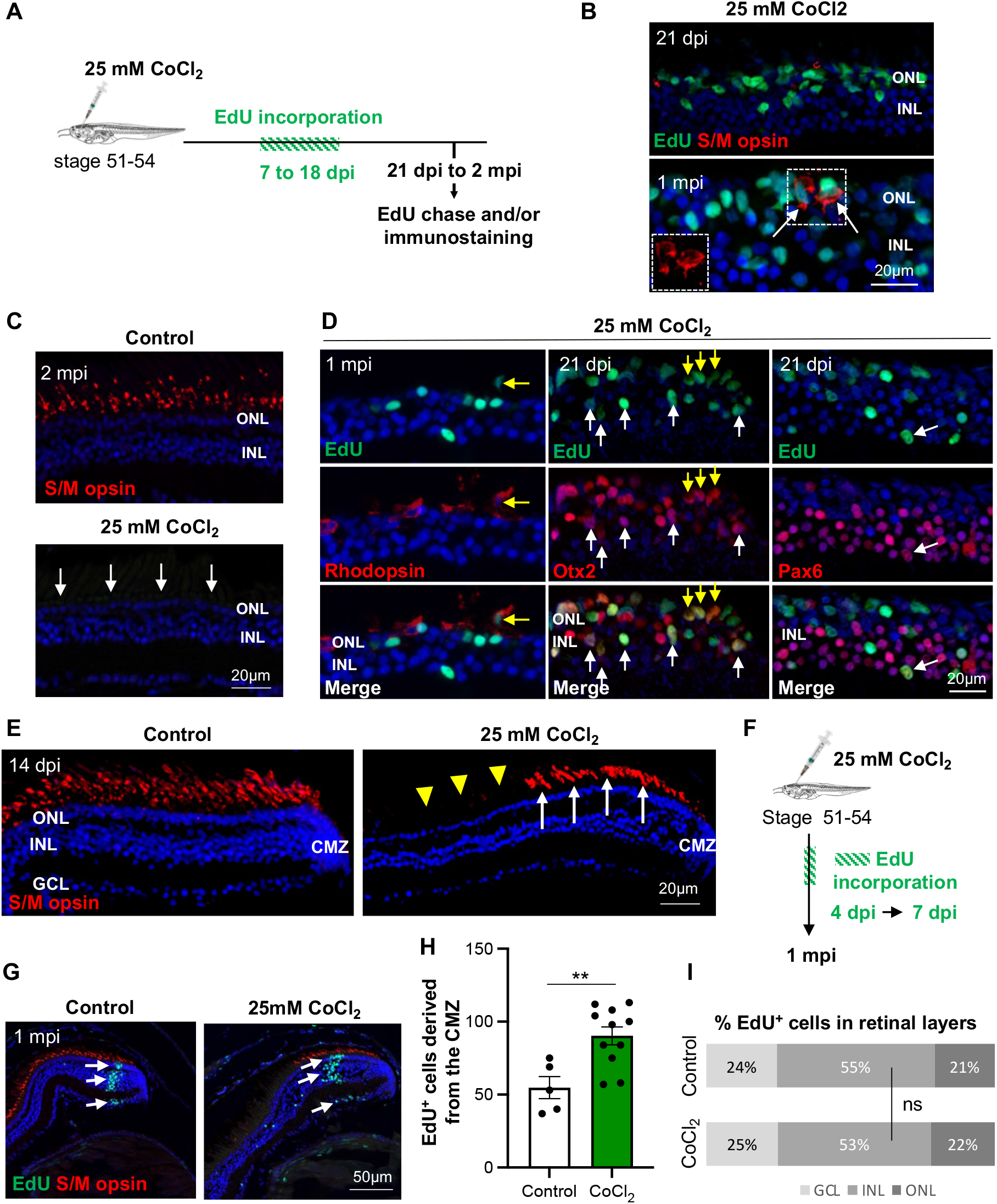
Absence of cone regeneration from Müller glial cells but CMZ-dependent cell replenishment, following a 25 mM CoCl_2_ intraocular injection in *Xenopus laevis*. **(A)** Outline of the experimental procedure used in (B-D). Stage 51-54 *Xenopus laevis* tadpoles were intraocularly injected with a saline solution (control) or with 25 mM CoCl_2_. They were then exposed to EdU from 7 to 18 dpi and finally processed for EdU staining and immunolabelling at 21 dpi, 1 mpi or 2 mpi, as indicated. **(B)** Representative images of retinal sections from CoCl_2_-injected tadpoles, co-labelled for EdU and S/M opsin at 21 dpi or 1 mpi. Arrows point to double-positive cells. S/M opsin staining alone is shown in the inset. **(C)** S/M opsin immunolabelling on retinal sections at 2 mpi. Arrows highlight the complete absence of staining in CoCl_2_-injected retinas. **(D)** Representative images of retinal sections from CoCl_2_-injected tadpoles, co-labelled for EdU and either Otx2, Pax6 (21 dpi) or Rhodopsin (1 mpi). White and yellow arrows point to double-positive cells in the INL and ONL, respectively. **(E)** Stage 51-54 tadpoles were intraocularly injected with a saline solution (control) or with 25 mM CoCl_2_. They were then processed for S/M opsin immunolabelling on retinal sections at 14 dpi. Shown here are representative images of retinal sections at the level of the CMZ. White arrows point to newborn cones produced by the CMZ. Yellow arrowheads indicate the absence of cones more centrally. **(F)** Outline of the experimental procedure used in (G-I). Stage 51-54 *Xenopus laevis* tadpoles were intraocularly injected with a saline solution (control) or with 25 mM CoCl_2_. They were then exposed to EdU from 4 to 7 dpi and finally processed at 1 mpi for EdU staining and immunolabelling. **(G)** Representative images of retinal sections stained for EdU and S/M opsin at 1 mpi. Arrows point to CMZ-derived newborn neurons (EdU^+^) in the different retinal layers. **(H)** Quantification of CMZ-derived EdU-positive cells. **(I)** Distribution of CMZ-derived EdU-positive cells within the three retinal layers. In all images, nuclei are counterstained with Hoechst. In (H), data are represented as mean ± SEM, and each point represents one retina (D). ns: not significant, **p < 0.01 (Mann-Whitney tests). GCL: Ganglion cell layer, INL: Inner nuclear layer, ONL: Outer nuclear layer, CMZ: Ciliary marginal zone.

Despite the virtual absence of S/M opsin staining within the central retina, labelled cells could be observed in the peripheral region at 14 dpi, likely corresponding to new-born cones produced by the CMZ (Fig. 7E). To confirm the occurrence of a CMZ-dependent regeneration process, we compared CMZ neuronal production of CoCl_2_-injected and control retinas through a EdU pulse-chase analysis (Fig. 7F). Consistent with the aforementioned enhanced proliferative activity of the CMZ, we found that more retinal cells were produced upon injury compared to the control situation (Fig. 7G, H). However, no difference in newborn cell distribution within the different retinal layers was observed, suggesting an absence of bias towards a particular cell fate (Fig. 7I). This shows that the CMZ do regenerate cells in this experimental paradigm, although not selectively those that have massively died.

### A high dose of CoCl_2_ triggers the formation of an ectopic mini retina derived from the RPE in *Xenopus laevis*

We next investigated whether the RPE proliferative bulge-like masses were able to differentiate. We found that they expanded over time, generating a massive extraocular tissue at 1 mpi, as assessed by 3D analysis on bleached tadpoles (Fig. 8A). This newly formed structure was highly variable in size and shape but presented with characteristics of a laminated retina. Analysis of retinal cell type revealed the presence of S/M opsin-positive cones (Fig. 8Bb). Of note, the mini retina-like structure was occasionally found inverted compared to the original retina lamination. In this case, cones could also be detected within the original retina, in the vicinity of the newly formed inverted one (Fig. 8Bc). The expression pattern of Rhodopsin, Otx2, Pax6, and Sox9 (a marker of Müller cells, Poché et al., 2008) further confirmed a layered organization as observed in a normal retina, with outer, inner, and ganglion cell layers (Fig. 8Ca-d). Besides, RPE65 labelling revealed that a new RPE had formed adjacent to the secondary photoreceptor layer (Fig. 8Ce). Finally, and unexpectedly, the newly formed retina occasionally contained a CMZ-like region, as assessed by a peripheral zone filled with EdU-positive cells (Fig. 8Cf).

**Figure 8.**
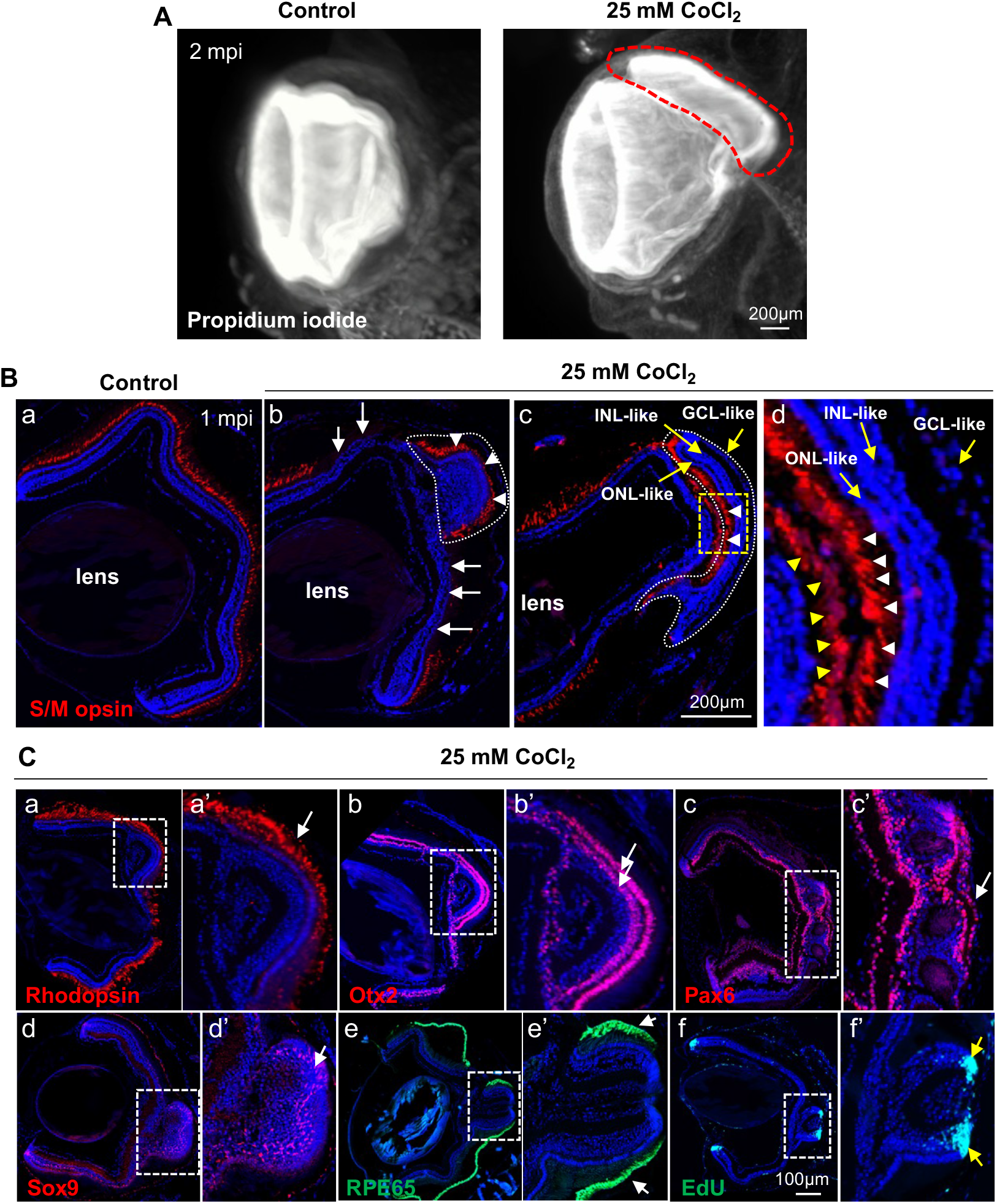
Formation of a layered retina-like structure from RPE, following a 25 mM CoCl_2_ intraocular injection in *Xenopus laevis*. **(A)** Stage 51-54 *X. laevis* tadpoles were intraocularly injected with a saline solution (control) or with 25 mM CoCl_2_. They were then processed at 2 mpi for tissue clearing and propidium iodide staining before being imaged by light-sheet microscopy. The red dotted line delineates the newly formed retina-like ectopic structure. **(B)** S/M opsin staining analysis at 1 mpi on retinal sections, following intraocular injection of a saline solution (control) or 25mM CoCl_2_. The dotted white lines delineate the RPE-derived ectopic retina-like structure. In panels (Bb) and (Bc), the retina-like structures show opposite orientations. (Bd) is an enlargement of the yellow dashed box in (Bc). White arrowheads point to new cones formed in the ectopic retina-like structures. White arrows in (Bb) highlight the absence of cone regeneration in the central retina when the ectopic structure has the same orientation as the original retina. In contrast, some regeneration occurs in contact with the ectopic structure, when this one is inverted (yellow arrowheads in (Bd)). Yellow arrows point to the three layers of the retina-like structure. **(C)** Immunodetection of different retinal cell markers or EdU labelling, as indicated, in CoCl_2_ injected retinas exhibiting an ectopic retina-like structure. Panels a’, b’, c’, d’, e’, and f’ are enlargements of the RPE-derived retina-like structures delineated by dotted boxes on the corresponding images. White arrows point to the labelled layers in these structures. Yellow arrows in f’ point to the CMZ-like regions. In all images, nuclei are counterstained with Hoechst. GCL: Ganglion cell layer, INL: Inner nuclear layer, ONL: Outer nuclear layer.

Together, these data highlight that in a degenerative context induced by 25 mM CoCl_2_ injection, three cellular sources contribute to the regeneration process: Müller glia mainly regenerate bipolar cells, the CMZ replenishes the peripheral retina, and RPE forms a retina-like structure facing the injured retina.

### A high dose of CoCl_2_ triggers Müller cell reactivation, RPE reprogramming, and enhanced CMZ proliferation in *Xenopus tropicalis*

*X. tropicalis* was so far described as using only its CMZ to regenerate its retina. Contrasting with *X. laevis*, no RPE reprogramming occurs following retinectomy (Miyake and Araki, 2014). In addition, we found that *X. tropicalis* Müller cells exhibit a very limited proliferative potential compared to *X. laevis* ones in a CRISPR/Cas9 model of rod degeneration (Parain et al., 2022). We thus questioned Müller and RPE cell response in our CoCl_2_ neurotoxic injury paradigm. As observed in *X. laevis*, intraocular injection of 25 mM CoCl_2_ in *X. tropicalis* tadpoles led to massive cell death in both the ONL and INL (Fig. 9A, B). Likewise, we found the same photoreceptor degenerative phenotype, with rods being less severely impacted than cones, which virtually disappeared from the central retina by 14 dpi onwards (Fig. 9C). Unlike their behaviour following specific rod degeneration (Parain et al., 2022), we surprisingly found that Müller cells efficiently re-entered into the cell cycle at 7 dpi, as inferred from PCNA and Yap co-labelling experiments (Fig. 9D, E and G-H). Of note, the average number of PCNA-positive cells per section was very similar to the one observed in *X. laevis* retinas (138 ± 6 *versus* 141 ± 12 at 7 dpi, respectively). The proliferative activity of the CMZ was also increased in these CoCl_2_-injected tadpoles compared to controls (Fig. 9D, F). Finally, RPE examination at 7 dpi revealed, similarly to *X. laevis*, morphological defects, as well as regions of proliferation associated with depigmentation (Fig. 9I). By 14 dpi, retina-like structures facing the degenerative retina and exhibiting cone labelling could occasionally be found (Fig. 9J). These data thus demonstrate for the first time that *X. tropicalis* can not only mobilize CMZ cells but also Müller glia and RPE, all of them behaving as in *X. laevis* to a CoCl_2_-driven neurotoxic retinal injury.

**Figure 9.**
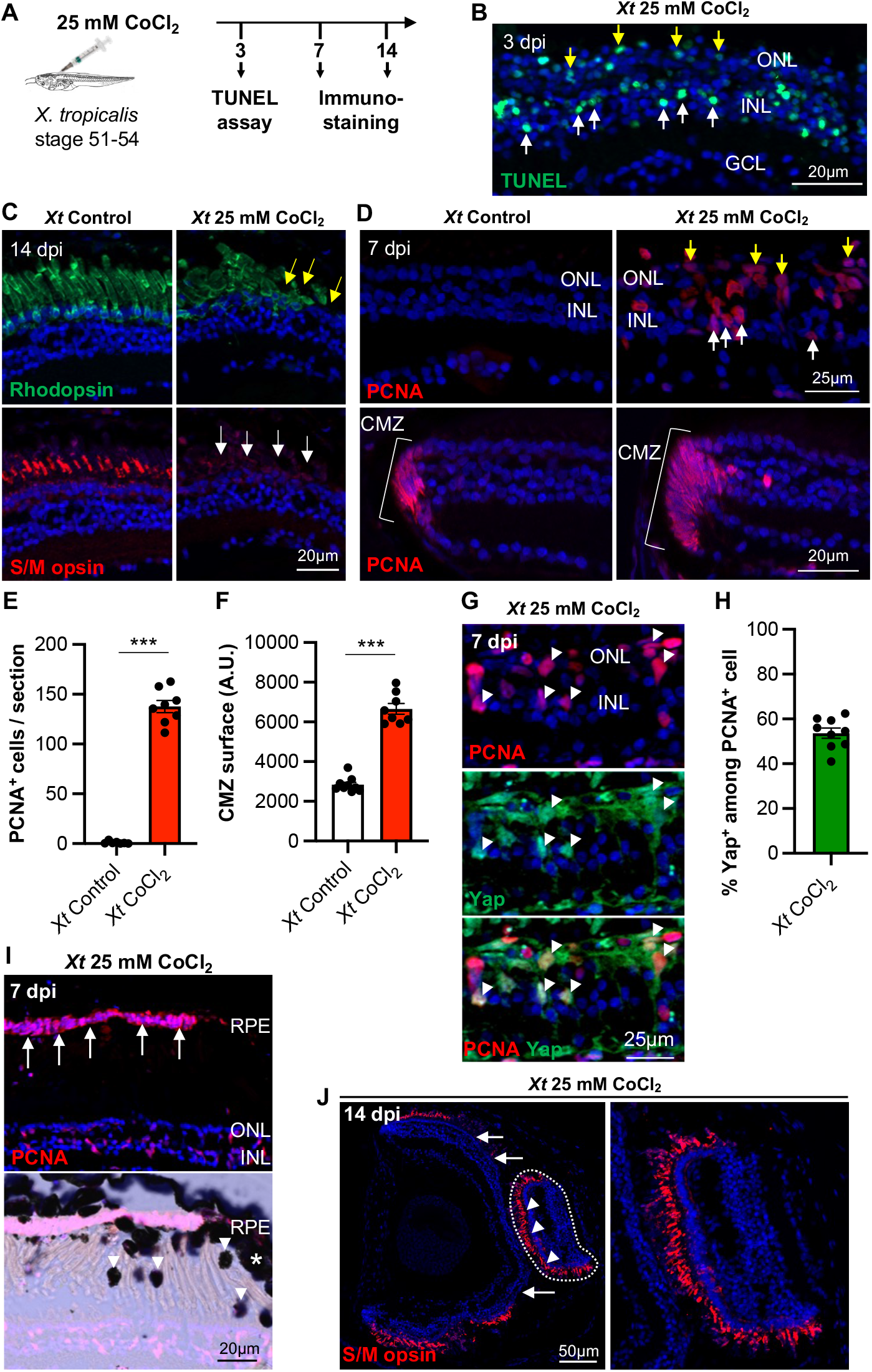
Mobilization of Müller glia, CMZ cells, and RPE in *Xenopus tropicalis*, following a 25 mM CoCl_2_ intraocular injection. **(A)** Outline of the experimental procedure used in (B-J). Stage 51-54 *Xenopus tropicalis* (*Xt*) tadpoles were intraocularly injected with a saline solution (control) or with 25 mM CoCl_2_. They were then processed at different time points, as indicated, for TUNEL assay or immunolabelling. **(B)** Representative image of retinal sections from CoCl_2_-injected tadpoles, stained with TUNEL at 3 dpi. White and yellow arrows point to positive cells in the INL and ONL, respectively. **(C)** Representative images of retinal sections stained for Rhodopsin or S/M opsin at 14 dpi. Yellow arrows point to defects in rod outer segment morphology, while white arrows highlight the absence of cones. **(D)** Representative images of retinal sections stained for PCNA at 7 dpi. The upper and lower panels show the central and peripheral regions of the retina, respectively. White and yellow arrows point to positive cells in the INL and ONL, respectively. **(E, F)** Quantification of PCNA-positive cells in the central retina (E) and CMZ surface (F) at 7 dpi. **(G)** Representative images of retinal sections co-labelled for PCNA and Yap at 7 dpi. Arrowheads point to double-positive cells. **(H)** Quantification of Yap^+^ cells among PCNA-labelled ones at 7 dpi. **(I)** Representative images (brightfield and PCNA labelling) of retinal sections from CoCl_2_-injected tadpoles at 7 dpi, showing proliferative RPE cells (arrows). On the brightfield image, arrowheads point to pellets of RPE, while asterisks indicate RPE thickening. **(J)** Representative images of retinal sections from CoCl_2_-injected tadpoles, stained for S/M opsin at 14 dpi. The dotted line delineates the RPE-derived ectopic retina-like structure (enlarged in the right panel), where new cones were formed (arrowheads). White arrows highlight the absence of cone regeneration in the central retina. In all images, nuclei are counterstained with Hoechst. In graphs, data are represented as mean ± SEM, and each point represents one retina. ***p < 0.001 (Mann-Whitney tests).

## DISCUSSION

Whether and how the extent of retinal damage or the nature by itself of the injury determines the regenerative modality remains an open question. We here characterized a new degenerative model in *Xenopus*, based on CoCl_2_-dependent neurotoxic damage, and showed that three different cellular sources could concomitantly be recruited for retinal repair. The advantage of this model is that CoCl_2_ affects different cell types in a dose-dependent manner. It thereby allows adjusting the extent of cell death. Although only Müller cells responded to the lowest concentration, a higher dose triggered a proliferative response of CMZ and RPE cells as well. Interestingly too, our data highlighted different neurogenic potentials of these cellular sources. In particular, we uncovered (*i*) the inability of *Xenopus* Müller glia to regenerate cones in this injury paradigm, although it is the most affected cell type, and (*ii*) the potential of RPE to reprogram and self-organize into a mini retina. Finally, this CoCl_2_-induced degenerative model also unveiled for the first time that *X. tropicalis* is able to reactivate its Müller cells and undergo RPE reprogramming, and not only to mobilize its CMZ, as previously hypothesized based on data from other injury models.

Modulating environmental oxygen levels has been shown to influence the rate of photoreceptor cell death in animal models of retinal degeneration (Yu and Cringle, 2005). Although cobalt toxicity is far from being understood, it has been determined that CoCl_2_ stabilizes the hypoxia-inducible factor (HIF) and thus mimics hypoxia, inducing in particular oxidative stress (Cervellati et al., 2014; Muñoz-Sánchez and Chánez-Cárdenas, 2019). Its ability to induce photoreceptor degeneration has been demonstrated *in vivo* in mice and rats (Hara et al., 2006), in monkeys (Shirai et al., 2016), and in zebrafish (Medrano et al., 2018). We found in *Xenopus* that cones undergo a more severe degenerative process than rods, as previously reported in zebrafish. The reasons underlying such different susceptibilities deserve further investigation. Interestingly, it has been shown that inducing hypoxia in RPE cells is sufficient to induce photoreceptor degeneration by reducing nutrient availability for the sensory retina (Kurihara et al., 2016). Photoreceptor cell death in our CoCl_2_ model may thus also be an indirect consequence of RPE metabolic stress.

CoCl_2_ injection is the first experimental paradigm leading to a virtual absence of cone photoreceptors in *Xenopus*. Although we previously found that Müller cells are able to produce new rods following their targeted ablation, we here found no sign of Müller glia-dependent cone regeneration. Interestingly, this contrasts with the zebrafish situation, where Müller cells do regenerate cones following CoCl_2_ treatment (Medrano et al., 2018). The reasons underlying such interspecies differences clearly require further investigation. The CoCl_2_ dose used might be involved. Indeed, the doses we had to inject were higher than those used in zebrafish to get similar phenotypes. It is thus possible that CoCl_2_ might have adverse effects on neurogenesis above a certain concentration threshold. Besides, such a need for higher CoCl_2_ concentrations suggests a lower sensitivity of *Xenopus* retinal cells to neurotoxicity. Overall, this novel *Xenopus* injury paradigm mimics the natural incapacity of human Müller cells to regenerate photoreceptors. As the ratio of cones to rods is 1:1 in *Xenopus* compared to 1:200 in mice, it thus represents a unique amphibian model for human age-related macular degeneration (AMD) and other cone-dystrophies. It should thereby offer the possibility to screen, in tadpoles and at a large scale, for molecules capable of driving Müller cell-derived progenitors towards a cone fate.

The ability of the RPE to undergo cellular reprogramming and regenerate retinal tissue has previously been reported after surgical removal of the entire retina in urodele and anuran amphibians, as well as in embryonic chicks (Araki, 2007; Barbosa-Sabanero et al., 2012; Chiba, 2014; Luz-Madrigal et al., 2014; Yoshii et al., 2007). Here again, interspecies differences have been identified, in particular between newt and *Xenopus*. Indeed, newt RPE cells undergo depigmentation, proliferate and regenerate a neural retina, all of it on-site on-site (Chiba, 2014). In contrast, *Xenopus* ones do not transdifferentiate inside their original location (Yoshii et al., 2007). Instead, a subpopulation detaches from the epithelium and migrates through the vitreous onto the retinal vascular membrane (RVM), where it forms a new neuroepithelium. The persistence of the RVM after retinectomy is thus assumed to be a prerequisite for regeneration to occur (Yoshii et al., 2007). In our CoCl_2_ injury model, we demonstrate the ability of the *Xenopus* RPE to reprogram and form a new retina locally, without any interaction with the RVM, suggesting the implementation of other inductive mechanisms.

In previous injury models that we developed, including (1) a mechanical needle poke injury (Langhe et al., 2017), (2) a transgenic line allowing for conditional rod ablation (Chesneau et al., 2018; Langhe et al., 2017), or (3) a CRISPR/Cas9-mediated *rhodopsin* knock-out (Parain et al., 2022), we never observed any ectopic retinal structures, as those appearing following 25 mM CoCl_2_ injection. The emergence of such mini retinas suggests that an eye field has formed from RPE cells, followed by self-driven retinal morphogenesis. Such *in vivo* organoid formation is reminiscent of the ectopic structures obtained following ectopic expression of eye-field transcription factors (Zuber et al., 2003). This phenomenon seems to depend on the injury paradigm. It also likely depends on the severity of the injury since lower doses of CoCl_2_ did not trigger RPE reprogramming, despite the loss of cones. Several hypotheses could account for the peculiar RPE cell response observed with 25 mM CoCl_2_. First, only this model, among the aforementioned ones, triggers massive apoptosis in both the ONL and INL. Compared with contexts where either cones or rods are selectively depleted, an extension of cell death to several cell types might trigger the release of specific key signals, and/or allow released signals to reach a threshold concentration sufficient for RPE mobilization. Second, RPE homeostasis was clearly perturbed upon 25 mM CoCl_2_ injection. It is possible that the stress caused by a hypoxic environment (possibly including enhanced oxidative stress), is a trigger for RPE reprogramming. Along this line, hypoxia was shown to enhance the efficiency of iPSC production and to cause the reprogramming of different differentiated cell types (Nakamura et al., 2021). Finally, CoCl_2_ injury causes inflammatory and angiogenic responses in zebrafish (Medrano et al., 2018). The resulting microenvironment may also be another trigger of RPE cell behaviour. This hypothesis is supported by the essential role played by the choroid, the vascular layer of the eye, in RPE-dependent regeneration (Ikegami et al., 2001; Kuriyama et al., 2009; Yoshii et al., 2007).

We recently reported in a model of selective rod degeneration, that *X. tropicalis* Müller cell proliferative potential was extremely limited compared to that of *X. laevis* (Parain et al., 2022). We found however that they actively re-enter into the cell cycle upon CoCl_2_-mediated injury. This demonstrates that *X. tropicalis* Müller glial cells are intrinsically able to get reactivated following retinal cell degeneration, although it likely requires different conditions than in *X. laevis*. Here again, the massive amount of apoptotic cells, the hypoxic-like environment, and the inflammatory and angiogenic responses, could all be factors underlying the differential behaviour of Müller cells in the CoCl_2_ model compared to a rod cell ablation context. Another difference between *X. laevis* and *X. tropicalis* is the RPE-dependent regeneration occurring upon complete removal of the retina. In this injury paradigm, only *X. laevis* exhibits this ability (Araki, 2007; Miyake and Araki, 2014; Yoshii et al., 2007). Why *X. tropicalis* RPE does not respond to retinectomy but does respond to CoCl_2_-induced retinal damage clearly deserves further investigation. Overall, such diversity of repair responses highlights the need to characterize how pathological microenvironments shape specific behaviours of different cellular sources.

## EXPERIMENTAL PROCEDURES

### Animals

*Xenopus laevis* and *Xenopus tropicalis* tadpoles were obtained by conventional procedures of *in vitro* or natural fertilization, and staged according to (Zahn et al., 2022). *Tg(her4:eGFP)* tadpoles were generated using the simplified REMI method (Chesneau et al., 2008; Sparrow et al., 2000), which consists only of co-injecting non-permeabilized sperm with the transgene. The transgene, designed by (Yeo et al., 2007), was previously shown to drive GFP expression in radial glial cells of the zebrafish brain (Galant et al., 2016). It was kindly provided by Laure Bally-Cuif’s laboratory.

### Intraocular injections

Stage 51-54 tadpoles were anesthetized in a solution of 0.005% benzocaine (Sigma) and then placed on a wet tissue in a petri dish. Intraocular injections were performed under a stereoscopic microscope using a Picospritzer III. Each eye received 2-3 30 nL-drops of CoCl_2_ (Sigma) at the indicated dose, or a saline solution as a control (0.1X MBS). Tadpoles were then put back in a tank filled with fresh water and monitored until they were awakened.

### Immunofluorescence, H&E staining

Tadpoles were deeply anesthetized in 0.01% benzocaine, fixed in 4% paraformaldehyde for 2 hours at room temperature and embedded in paraffin. They were then sectioned (11 μm) with a Microm HM 340E microtome (Thermo Fisher Scientific).

Immunostaining was performed using standard procedure. Briefly, after xylene dewaxing, antigen retrieval was performed by boiling the sections in 10 mM sodium citrate and 0.05% Tween20 (Sigma) for 10 min. After a 1-hour blocking step in 10% goat serum (Life technologies), 5% BSA (Sigma) and 0.2% Tween20, slides were incubated with primary antibodies (Supplementary Table 1) overnight at 4°C, and then for 2 hours at room temperature with secondary antibodies (Alexa Fluor; Thermo Fisher Scientific). In some instances, bleaching of RPE was necessary. To this end, sections were incubated for about 15 min at 55°C in 10% hydrogen peroxide, prior to unmasking. Cell nuclei were counterstained with Hoechst (10 μg/mL, Sigma) and coverslips were mounted using Fluorsave Reagent (Millipore).

Hematoxylin and eosin (H&E, RAL Diagnostics) staining was performed according to standard procedures.

### EdU labelling

Tadpoles were immersed in a solution containing 1 mM EdU (5-ethynyl-2’-deoxyuridine; Invitrogen) before fixation. The solution was replaced every other day. EdU labelling was performed using Click-iT® Plus EdU Imaging Kits (Invitrogen).

### Tissue clearing for light sheet microscopy

Tissue clearing was performed on albinos *Xenopus laevis* tadpoles as described in (Matsumoto et al., 2019). Nuclear staining was performed by immersion for 3 days in a 30 μg/mL propidium iodide solution.

### Microscopy

Fluorescence and brightfield images were obtained with an Axio Imager M2 microscope (Zeiss). They were then processed using Zen (Zeiss, Germany), Image J (Schneider et al., 2012), and Photoshop CS5 (Adobe) softwares. Three-dimensional images following tissue clearing were acquired with a light-sheet microscope UltraMicroscope II (Miltenyi Biotec).

### Quantification, and statistical analysis

Quantification of TUNEL-, EdU-, Otx2-, Pax6-, and PCNA-positive cells was performed by manual counting within the neural retina (excluding cells from the CMZ). In situations where the number of apoptotic cells and apoptotic bodies were too high for reliable manual counting, we rather measured the staining area using Image J software, after setting an optimal intensity threshold. The same approach was used to quantify S/M opsin staining. CMZ area was measured on PCNA- or EdU-labelled retinas using Photoshop CS5 software. To estimate the percentage of co-labelled cells, quantifications were performed in the center of the retina, within a rectangle of 200 × 100 μm. Depending on the experiments, 3 to 8 sections per retina were analyzed. An average number was then calculated for each retina and represented by one dot on the dot plots. All experiments were performed at least in duplicate. Figures show the results from one representative experiment. Statistical analyses were performed using the non-parametric Mann-Whitney test. Statistical significance is: *p < 0.05; **p < 0.01; ***p < 0.001; ns: not significant.

### Ethics statement

All animal care and experimentation were conducted in accordance with institutional guidelines, under the institutional licence C 91-471-102. The study protocol was approved by the institutional animal care committee CEEA #59 and received authorization by the Direction Départementale de la Protection des Populations under the reference number APAFIS #21474-2019071210549691 v2 and #32589-2021072719047904 v4.

## ACKNOWLEDGEMENTS

We are thankful to L. Bally-Cuif for the gift of the *Her4:GFP* construct. We are grateful to A. Donval for her help with *in vitro* fertilizations. This research was supported by grants to M.P. from the Association Retina France, Fondation Maladies Rares, Association du syndrome de Bardet-Biedl (BBS), Fondation de France, and UNADEV (Union Nationale des Aveugles et Déficients Visuels) in partnership with ITMO NNP (Institut Thématique Multi-Organisme Neurosciences, sciences cognitives, neurologie, psychiatrie) / AVIESAN (alliance nationale pour les sciences de la vie et de la santé).

## SUPPLEMENTARY FIGURE LEGENDS

**Figure S1.**
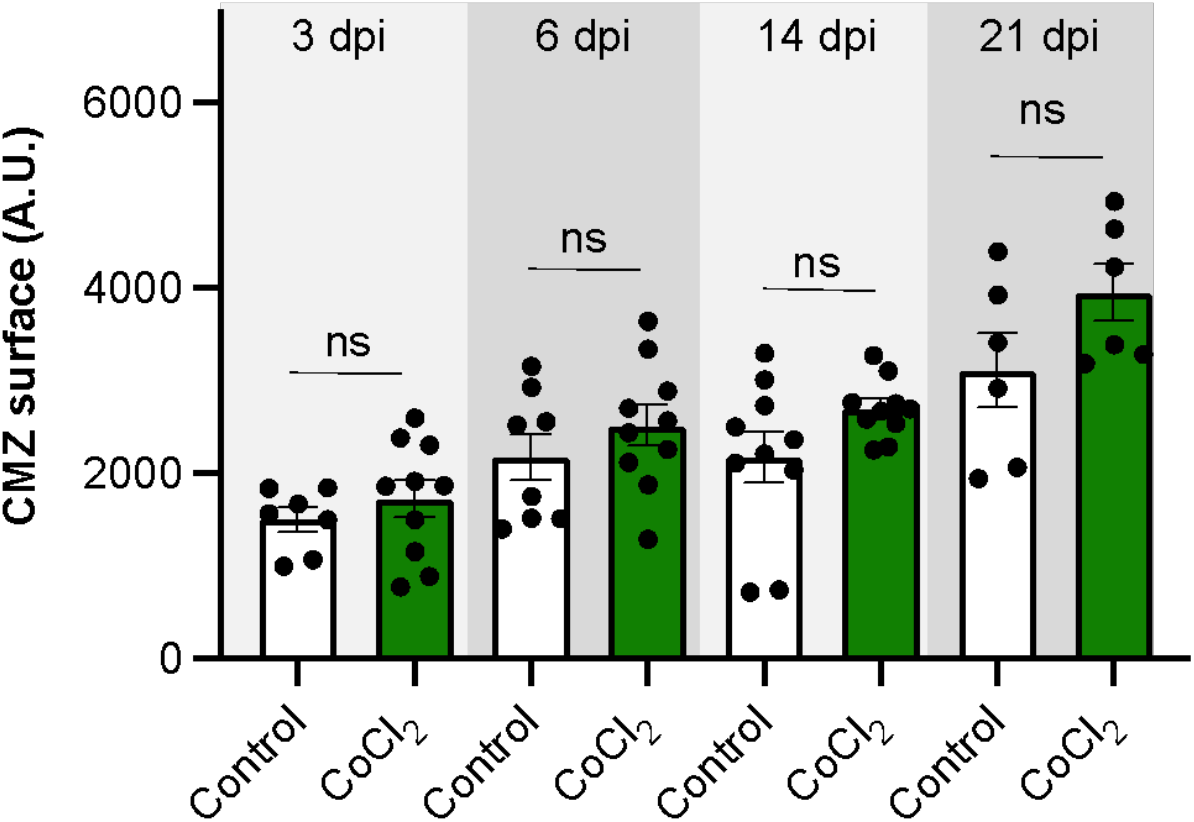
Absence of CMZ cell proliferative response, following intraocular injection of 10 mM CoCl_2_ in *Xenopus laevis*. Quantification of CMZ surface (as assayed by EdU labelling) at 3, 6, 14, or 21 dpi in stage 51-54 *X. laevis* tadpoles, following injection of a saline solution (control) or 10 mM CoCl_2_. Data are represented as mean ± SEM, and each point represents one retina. ns: not significant (Mann-Whitney tests).

**Supplementary Table 1.**
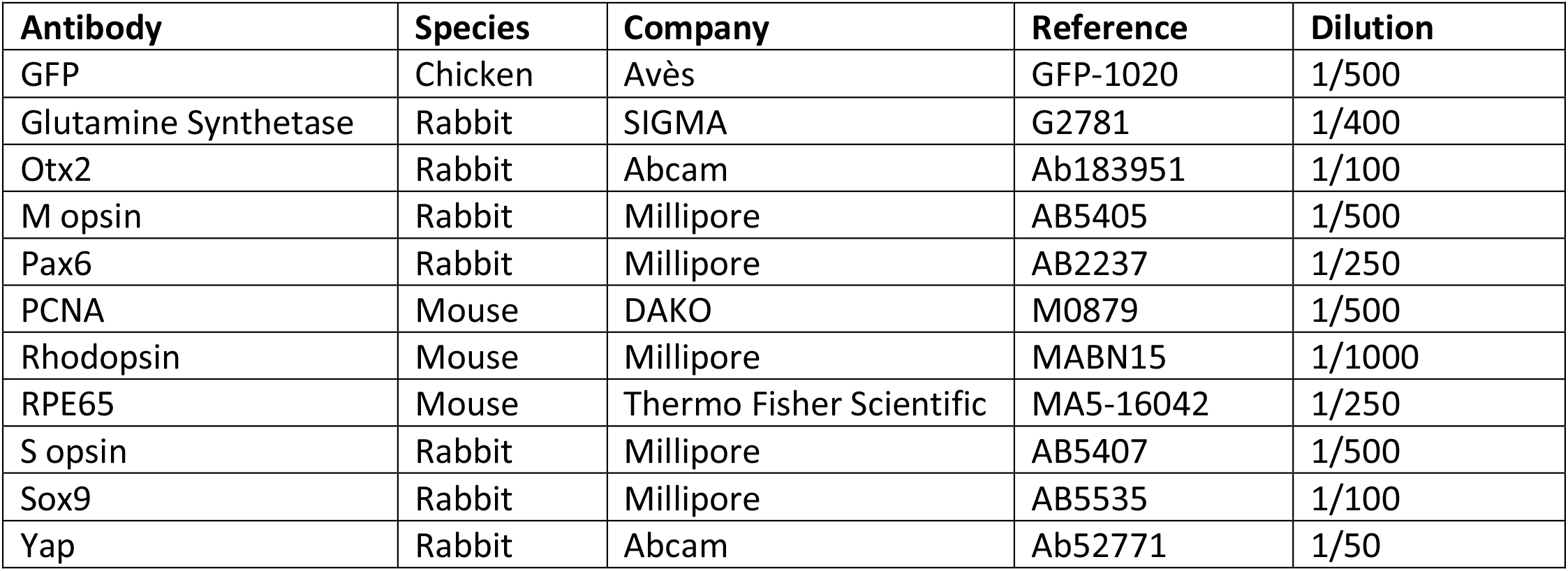

| **Antibody** | **Species** | **Company** | **Reference** | **Dilution** |
| --- | --- | --- | --- | --- |
| GFP | Chicken | Avès | GFP-1020 | 1/500 |
| Glutamine Synthetase | Rabbit | SIGMA | G2781 | 1/400 |
| Otx2 | Rabbit | Abcam | Ab183951 | 1/100 |
| M opsin | Rabbit | Millipore | AB5405 | 1/500 |
| Pax6 | Rabbit | Millipore | AB2237 | 1/250 |
| PCNA | Mouse | DAKO | M0879 | 1/500 |
| Rhodopsin | Mouse | Millipore | MABN15 | 1/1000 |
| RPE65 | Mouse | Thermo Fisher Scientific | MA5-16042 | 1/250 |
| S opsin | Rabbit | Millipore | AB5407 | 1/500 |
| Sox9 | Rabbit | Millipore | AB5535 | 1/100 |
| Yap | Rabbit | Abcam | Ab52771 | 1/50 |

